# Classifying Aging as a Disease in the context of ICD-11

**DOI:** 10.1101/024877

**Authors:** Alex Zhavoronkov, Bhupinder Bhullar

## Abstract

Aging is a complex continuous multifactorial process leading to loss of function and crystalizing into the many age-related diseases. Here, we explore the arguments for classifying aging as a disease in the context of the upcoming World Health Organization’s 11th International Statistical Classification of Diseases and Related Health Problems (ICD-11), expected to be finalized in 2018. We hypothesize that classifying aging as a disease will result in new approaches and business models for addressing aging as a treatable condition, which will lead to both economic and healthcare benefits for all stakeholders. Classification of aging as a disease may lead to more efficient allocation of resources by enabling funding bodies and other stakeholders to use quality-adjusted life years (QALYs) and healthy-years equivalent (HYE) as metrics when evaluating both research and clinical programs. We propose forming a Task Force to interface the WHO in order to develop a multidisciplinary framework for classifying aging as a disease.

## Introduction

The recognition of a condition or a chronic process as a disease is an important milestone for the pharmaceutical industry, academic community, healthcare and insurance companies, policy makers, and individual, as the presence of a condition in disease nomenclature and classification greatly impacts the way it is treated, researched and reimbursed. However, achieving a satisfactory definition of disease is challenging, primarily due to the vague definitions of the state of health and disease. Here, we explore the potential benefits of recognizing aging as a disease in the context of current socioeconomic challenges and recent biomedical advances.

## Brief History of Disease Classification

The concept of disease classification has existed for centuries, with professional classification dating back to 15^th^ century Italy, but international classification systems were not established until relatively recently. One of the first major strides was made by William Cullen, who provided a classified list of diseases in his Nosolagae Methodicae synopsis in 1769. The first authoritative sources of disease terminology were then developed in 19th century England (Robb-smith 1969; Moriyama et al., 2011), followed by the nomenclature of diseases by the American Medical Association (National Conference on Nomenclature of Disease, 1933; American Medical Association, 1961). Real progress toward the modern classification system, however, began when efforts were internationalized. The first international nomenclature and disease classification was created and maintained by the International Statistical Institute (ISI) and is generally referred to as ICD-1, as it was developed at the first international conference to revise the International Classification of Causes of Death in 1900. Since 1948, the World Health Organization (WHO) has assumed the responsibility of maintaining, preparing and publishing the revisions of the ICD. The 10th revision of the ICD, referred as ICD-10, was first published in 1992 (World Health Organization, 1992), and the 11th revision (ICD-11) is expected to be released in 2018 (http://www.who.int/classifications/icd/revision/timeline/en/). Thus, as this date approaches there is a sense of urgency among all stakeholders to codify classification proposals before the window of opportunity closes for another decade or longer (World Health Organization, 2006).

In its current definition, published in 1948, WHO describes health as a “state of complete physical, mental and social well-being, not merely the absence of infirmity” (World Health Organization, 1948). This definition is restrictive and there have been an increasing number of calls to change it (Saracci, 1997;Bircher, 2005;Huber et al., 2011). The criteria for disease, on the other hand, have historically changed over time. This is partly due to increasing health expectations or changes in diagnostic abilities, but mostly due to a combination of social and economic reasons (Scully, 2004). It was not until the 6th revision of ICD (ICD-6) in 1949 that mental disorders were first classified as diseases (Katschnig, 2010). On the other extreme, homosexuality was classified as a disease until 1974 by the American Psychiatric Association (APA) (Reznek, 1987), an organization which frequently disagrees with the ICD on disease classification. Since the completion of ICD-10, debates in biomedical literature have been primarily focused on psychiatric disorders (First et al., 2015). A recent PubMed search using “ICD-11” since 1992 produced 333 results, while “’ICD-11’ NOT mental NOT psychiatric” produced only 61 results.

While mental health issues continue to stay at the forefront, another major socioeconomic problem has surfaced and is here to stay. Over the past decade, there has been an enormous shift in the percentage of the world population that is elderly, and age demographics are projected to continue to change dramatically in this direction in the next few decades. The socioeconomic burden of this shift cannot be understated. Without advances in the ability to slow the aging process, extend the healthy life span, and prevent the associated diseases, it will be difficult to support the large numbers of those no longer enrolled in the workforce and instead enrolled in costly long term health care.

Fortunately, the ability to track multiple underlying causes of death has provided a more granular view on the causes of death in old age and in turn has facilitated better disease association and classification (Moriyama et al., 2011). Our understanding of aging mechanisms is constantly evolving. Similarly, disease classification is an evolving process and advances in our understanding of aging may enable the classification of aging as a disease in the upcoming revision of the ICD. To do so would have multiple benefits, garner more attention on the topic, and ultimately boost progress in this active area of biomedical research.

## Biology of aging as a disease

Aging is a complex multifactorial process leading to loss of function, multiple diseases, and ultimately death. There are many theories explaining the origin of the overall process (Lopez-Otin et al., 2013;Deleidi et al., 2015), cause and effect relationships between different processes and systems, including aging of the immune system (Montecino-Rodriguez et al., 2013), inflammation (Bruunsgaard et al., 2001;Sarkar and Fisher, 2006;Franceschi et al., 2007;Michaud et al., 2013), fibrosis (Cieslik et al., 2011;Kapetanaki et al., 2013), mineralization of connective tissue (Shindyapina et al., 2014), cellular senescence (van Deursen, 2014), wear and tear, and many others. In addition, many genetic and epigenetic changes implicated in aging and longevity are associated with aging in model organisms (Lombard et al., 2005;Moskalev et al., 2014). Even though their role and action in human aging are uncertain, many of these changes are also associated with human diseases (Kennedy et al., 2015; Aguilar-Olivos et al., 2015;De Rosa et al., 2015;Helling and Yang, 2015;Lardenoije et al., 2015;Renauer et al., 2015). Recent discoveries showing that mechanisms involved in cancer are also associated with the aging process have led to multiple proposals to prevent cancer and other age-related diseases using drugs that increase lifespan in model organisms (Blagosklonny, 2013;Zhavoronkov et al., 2014).

Surprisingly, aging shares many characteristics with Human Immunodeficiency Virus (HIV), as while having a specific ICD-10 code, HIV is the cause of many fatal diseases associated with a broad range of ICD-10 codes. However, unlike HIV, classifying aging as a disease is difficult because of the absence of a clear set of aging biomarkers and the uncertainty of the time of transition from loss of function to disease. There is some progress in this area; recent population statistics and studies clearly demonstrate the relationship of aging and increased risk of multiple diseases (Marengoni et al., 2011;Federal Interagency Forum on Aging-Related Statistics, 2012;Salive, 2013), and there are several frailty indexes that measure the rate of physical decline and sets of aging biomarkers (Moreno-Villanueva et al.;Horvath, 2013). However, the transition from aging to disease remains unclear both at the individual and population levels (Figure 1).

**Figure 1:**
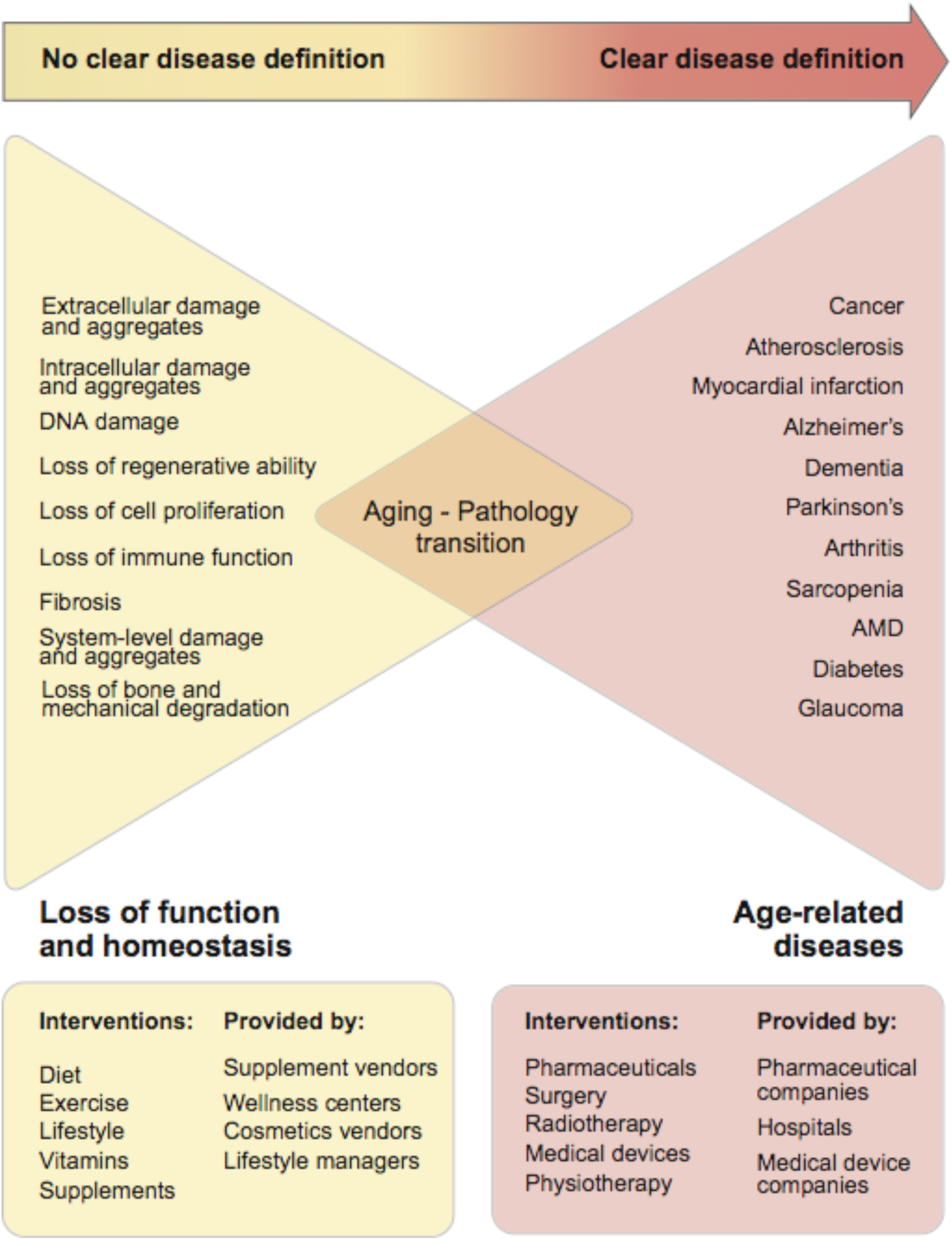
Gradual transition from the loss of function and homeostasis due to aging into age-related pathologies. It is clear that with time the combination of many age-related processes becomes more granular and crystallizes into specific diseases. However, it is unclear when aging starts and when the transition between aging and pathology occurs.

## Public opinion and stakeholders’ views on aging as a disease

At the start of the twentieth century, most causes of death in advanced age were attributed to “old age” and “natural causes” (Moriyama et al., 2011). However, over the course of the twentieth century, the understanding of aging and disease changed (Shock et al., 1984;Bulterijs et al., 2015;Faragher, 2015), with many age-related diseases and causes of death being clearly defined (Vellas et al., 1992). Over time, many chronic diseases of old age previously considered to be a part of normal aging including hypertension, rheumatoid arthritis, cardiomyopathy, and osteoporosis received classification codes (Bennett et al., 1956;force, 1980;WHO, 1994). Historically, “old age” was among the acceptable causes of death and was used on death certificates (Moriyama et al., 2011). Most countries now provide guidelines limiting the use of the term on medical certificates by age and issuing recommendations to avoid using only “old age” as a cause of death (Office for National Statistics, 2010), since “terms such as senescence, infirmity, old age, and advanced age have little value for public health or medical research” (Centers for Disease Control, 2004). The same arguments are commonly used in debates against classifying aging as a disease, as many stakeholders would prefer to have a more granular view into causes of death.

To illustrate these differences in opinion about aging, a recent survey in Finland assessed stakeholders’ perspectives on what constitutes a disease. The study surveyed 3,000 laypeople, 1,500 doctors, 1,500 nurses and 200 parliament members and asked them to rate 60 “states of being” by their perception of disease. And while most cancers like breast cancer (#1) and prostate cancer (#2) were clearly classified as diseases, wrinkles (#60), smoking (#59) and ageing (#58) were not classified as diseases (Tikkinen et al., 2012). Most of the doctors and nurses did not consider sarcopenia, age-related muscle loss (#31) as a disease. Surprisingly, obesity (#50), was not considered to be a disease by any of the stakeholder groups even though it is now classified as a disease with multiple disease codes (E65-E68) in endocrine, nutritional and metabolic diseases (E00-E90), representing a considerable part of chapter IV in ICD-10 (World Health Organization, 2011). The case of obesity provides clues that the opinion of multiple stakeholders including medical, general public, doctors and nurses may deviate from the WHO consensus for the purposes of disease classification, and other stakeholders may play a larger role in forming ICD revisions.

In another survey of pension funds, insurance companies, and employee benefits industries; only 11% of respondents felt that aging is a disease and less than 22% used simulations involving biomedical advances in forecasting life expectancy (Zhavoronkov, 2015). Considering the relatively conservative responses, these stakeholder groups may not be in favor of classifying aging as a disease in the ICD-11 revision.

The debate on whether aging should be classified as a disease is taking place primarily in academic circles (Ries, 1976;Glasser, 1986;Williams, 1987;Blumenthal, 1993;Callahan and Topinkova, 1998;Blumenthal, 2003;Faragher, 2015;Gems, 2015b) with no consensus opinion (Garber, 2008;Donmez and Guarente, 2010) and even some opposing views from biogerontologists (Ardeljan and Chan, 2013;Rattan, 2014). Even though some of these views aim at higher citation rating or other objectives, these opposing views do not propose concrete plans for addressing the aging problem at the level of WHO.

Prior history with many mental disorders, including autism (Lord and Jones, 2012), demonstrates that classifying a state of being as a disease leads to an increased attention to the subject, the development of more accurate diagnostic methods, and increased involvement of the pharmaceutical industry and policy makers. It also provides the basis for clinical trials.

## Strategies in moving forward

### Large scale studies in humans

There are many strategies that can be pursued to test the efficacy of geroprotectors in humans, including large scale publicly-funded supplement studies and studies designed to address a specific set of biomarkers of aging (Hefti and Bales, 2006;Le Couteur et al., 2012;Scott and DeFrancesco, 2015). The results from the Novartis RAD001 study in healthy elderly patients as a vaccine-potentiating agent (Mannick et al., 2014) elucidated the possibility of ameliorating immunosenescence in the elderly and raises the prospect of delaying the onset of age-related diseases.

### Aging biomarkers

Even though major advances have been made since the final ICD-10 meeting in tracking aging at all levels of organization (Sprott, 2010;Le Couteur et al., 2012;Hatse et al., 2014;Wu et al., 2015), there is no universal set of biomarkers and guidelines for measuring aging as a system. However, in order to successfully evaluate the effect of any drug that influences aging, it is essential have a measureable endpoint, such as biomarkers. Gerontologists have previously struggled to extrapolate biomarkers from animal models to humans (Butler et al., 2004). But with the advent of Big Data, it is now possible to track aging on the epigenetic level and measure accelerated aging in many diseases (Horvath et al., 2014;Horvath and Levine, 2015). There are also promising studies of transcriptomic (Nakamura et al., 2012;Dhahbi, 2014), telomere-length (Zhang et al., 2014;Shamir, 2015) and multi-variate (Sanders et al., 2014) blood-based biomarkers that may lead to minimally invasive diagnostic tests. It is also possible to track signalome-level biomarkers. Recent studies have shown that signaling pathways found in aging are comparable to and share many characteristics with Hutchinson-Gilford Progeria Syndrome (HGPS) (Aliper et al., 2015) and Age-related Macular Degeneration (AMD) (Makarev et al., 2014). It is also possible to use system-wide biomarkers like heart rate variability (HRV) as biomarkers of aging (Corino et al., 2007). There is a rapidly growing body of evidence that biomarkers of aging contributes and is very similar to the many age-related diseases on all levels of organization and it is possible to multiplex epigenetic, transcriptomic, proteomic, signalomic, metabolomic, metagenomic and phenotypic biomarkers to track the progression of aging as a disease.

Since the transition from age-associated processes to disease is unclear (Figure 1), in the absence of readily available and highly personalized biomarkers, one of the approaches to set the gold standard for health, is to select the age when peak performance is observed in the majority of the population. There are conflicting theories without convincing empirical evidence that the human skeleton stops growing at approximately the age of 20 (Nilsson et al., 2005). Peak age for athletic performance depends upon sports disciplines and is usually in the range of 20-30 years with younger age of peak performance in short distance races and gradually increasing age of peak performance in longer racing distances (Elmenshawy et al., 2015). In prenatal diagnostics, statistics for Down’s and other chromosomal abnormalities led to recommendations for invasive prenatal screening after 35, which is generally referred to as advanced maternal age, procedures that have gradually been replaced by non-invasive prenatal procedures. (Chiu et al., 2009;Nakata et al., 2010;Benn et al., 2013;Lo, 2013;Nepomnyashchaya et al., 2013;Twiss et al., 2014;Neufeld-Kaiser et al., 2015).

## Discussion

Many scientists throughout the world have argued that aging should be classified as a disease (Bulterijs et al., 2015;Gems, 2015a). The recent decision of the Food and Drug Administration (FDA) to test the potential of metformin in reducing risk of developing age-related conditions including cancer, heart disease and cognitive impairment resonated in the community, increased attention to the field, and suggested that aging may be an indication appropriate for clinical trials (Check Hayden, 2015). However, there are no visible global or national efforts to include aging in the ICD-11 classification as a disease. This is primarily due to the structure and clear separation of the supplement and wellness markets and disease treatment markets. Stakeholders in the wellness market are selling billions of dollars worth of goods and services and are not interested in increased regulation. Also, the structure of the supplement market, where large ingredient vendors supply product to compounders that in turn supply brand holders that sell through distribution networks, does not provide incentives or ability at any level to engage in discussions with policy makers.

As demonstrated in Figure 1, individual consumers of healthcare cannot formulate the demand for evidence-based medicine (pharmaceuticals) to combat aging and rely on the less-regulated wellness and alternative medicine industry. The fact that the supplement industry is prosperous, with a total estimated sale of $36.7 billion in the US in 2014, implies that there is consumer demand for anti-aging treatments (Nutrition Business Journal, 2015). Increasing regulation of the supplements and wellness industry may lead to more demand and a supply of evidence-based approaches to interventions in aging.

Increasing productive longevity in the developed countries is likely to reduce healthcare costs, boost employee productivity and lead to substantial economic growth (Zhavoronkov and Litovchenko, 2013), while the inability to increase productive longevity quickly may result in economic collapse (Zhavoronkov et al., 2012). Yet when it comes to aging, policy makers, the general public, and even pension funds lack the sense of urgency to address it as a curable disease (Zhavoronkov, 2015) even though there is a growing realization of the likely economic problems (Zhavoronkov, 2013). Even some of the most reputable demographers, gerontologists and biogerontologists express conflicting opinions on the subject, providing arguments against classifying aging as a disease. While these papers promote debate and increase their citation ratings, they are certainly not helpful for making the case for formal classification of aging as a disease or family of diseases. Formal classification of aging as a disease is likely to unite both scientists and medical practitioners in this effort.

There are also ethical considerations associated with classifying aging as a disease (Caplan, 2005). A large portion of the population may feel uncomfortable with the idea and resist being classified as disease carriers either at birth or after a specific age certain age while lacking clear representation of a disease. However, the recent inclusion of obesity in ICD-10 has shown that it not only helped to attract resources to research, but it also resulted in more frequent diagnosis of the condition. Since obesity has been classified as a disease, it is now easier for medical practitioners to convince patients to pursue healthier lifestyles and prescribe medication, even in cases where patients are comfortable with the condition.

## Recommendations for Implementation

### Form a task force

As part of the upcoming ICD-11, the International Association for the Study of Pain (IASP) Task Force was created to classify chronic pain as a disease (Treede et al., 2015), with a clear organizational structure to interact with the WHO to create a classification system of all manifestations of chronic pain, resulting in 7 categories: musculoskeletal pain, visceral pain, headache, neuropathic pain, post surgical and posttraumatic pain, cancer pain and primary pain. A similar task force should be created to interact with the WHO on the classification of aging.

### Develop an “ideal norm”

Despite the growing abundance of biomarkers of aging, classifying aging as a disease will be challenging due to the absence of the “ideal norm.” Despite significant effort from the academic and industry communities, sarcopenia is still not classified as a disease despite clear clinical and molecular representation and similarity with premature musculoskeletal aging and myotonic disorders (Cruz-Jentoft et al., 2010;Muscaritoli et al., 2010;Fielding et al., 2011;Morley et al., 2011;Mateos-Aierdi et al., 2015). One approach to address this challenge is to assume an “ideal” disease-free physiological state at a certain age, for example, 25 years of age, and develop a set of interventions to keep the patients as close to that state as possible. Considering the WHO definition of health, it may be possible to agree on the optimal set of biomarkers that would be characteristic to the “state of complete physical, mental and social well-being, not merely the absence of infirmity” and agree on the physiological threshold after which the net totality of deviation of these biomarkers from norm can be considered a disease.

### Set effective metrics

When performing cost-effectiveness analysis (CBA), economists often evaluate the outcomes of various programs in terms of the quality-adjusted life years (QALYs) and healthy-years equivalent (HYE) (Mehrez and Gafni, 1989). The most effective altruistic causes are now also evaluated using these metrics (Macaskill, 2015). Studies also demonstrate that each QALY can be valued at $24,777 to $428,286 depending on the method (Hirth et al., 2000). Many projects in aging research do not result in QALY and are currently not prioritized to maximize healthy lifespan (Zhavoronkov, 2013). Classifying aging as a disease would facilitate the development of programs and prioritize projects in aging research that maximize QALY and HYE today and in the future.

## Conclusion

There are definite benefits for many stakeholders in having aging classified as a disease, and the research community should consider uniting and working as a single voice to increase the chance of having aging classified as a disease by the WHO.

## Acknowledgements

We would like to thank Hantamalala Ralay Ranaivo for reviewing and editing this manuscript.

